# Hormonal factors are poor indicators of endometriosis pain

**DOI:** 10.64898/2026.09.29.755136

**Authors:** Adam J. Dourson, Rachel L. Cundiff-O’Sullivan, Juliet Mwirigi, Holly R. Hoefgen, Maggie Dwiggins, Elise Bardawil, Thomas J. Baranski, Benjamin P. Kay, Robert W. Gereau, Whitney T. Ross, Hadas Nahman-Averbuch

## Abstract

Endometriosis is a common yet understudied condition in which sex hormones play a key role in the etiology and pathophysiology of the disease. Sex hormones are dysregulated in endometriosis and are pharmacologically targeted to reduce symptoms including pain, but the relationships between endometriosis-related pain with sex hormone levels and their receptors are not clear. In 60 women previously diagnosed or undergoing laparoscopic surgery for the diagnosis of endometriosis, we collected patient-reported endometriosis/pelvic pain intensity in the last 30 days, pain interference scale, and the Endometriosis Health Profile 30 (EHP30) pain subscale. In a sub-sample of 38 participants, biopsies from control non-lesion pelvic peritoneum and endometrial lesion tissues were collected during surgery to analyze the tissue abundance of the primary receptors for estrogen, as well as androgen, progesterone, and prolactin receptors. Serum was collected to analyze systemic sex hormone levels including estradiol, prolactin, progesterone, and testosterone. Between tissues, we found elevated transcript levels for estrogen receptor alpha (*ESR1*) in lesions, but no difference in estrogen receptor beta (*ESR2*) transcript levels between tissue samples were found. Relating tissue receptor transcript levels to clinical pain revealed that only *ESR1* and *PGR* in lesions were negatively associated with endometriosis/pelvic pain intensity, however, following false discovery rate correction the association with ESR1 was no longer statistically significant. No other tissue or systemic factors were associated with clinical pain outcomes. Together, these data demonstrate unique alterations in receptor abundance in lesions yet select transcript levels only weakly relate to pain in endometriosis.

**One Sentence Summary:** Endometriosis-associated pain is largely unrelated to serum sex hormone levels and receptor expression levels in tissue.

## INTRODUCTION

Endometriosis affects approximately 200 million individuals worldwide with annual healthcare expenditures estimated at $69.4 billion (*1, 2*). It is defined by the presence of ectopic, endometrial-like lesions implanted outside the uterus, the extent and location of which define clinical staging of the disease. Chronic pelvic pain is the most common symptom of endometriosis, with 37% of patients reporting severe pain (*3–5*). Importantly, the clinically determined stage of endometriosis does not reliably correlate with the intensity of pelvic pain; for example, patients who report severe pain often present with few superficial lesions and are assigned disease stage I (*6–8*). Such disconnects underscore the need to better understand the factors driving endometriosis pain, which remains understudied in both mechanistic animal models and clinical cohorts (*4, 9–14*). A more comprehensive understanding of the factors related to pain associated with endometriosis is essential to optimize current treatments and to inform the development of novel, targeted therapies.

Despite multiple strategies to treat endometriosis, successful management of symptoms is inconsistent. Currently, the most common treatments to alleviate endometriosis pain are hormonal medications designed to suppress ovulation and menstruation by targeting the hypothalamic-pituitary-ovarian axis (*15–18*). A systematic review assessing treatments for endometriosis patients found that 5-59% of patients reported some pain remaining at the end of treatment while 11-19% reported no change in pain symptoms after treatment (*19*), reinforcing that a better understanding of underlying mechanisms is vital to improve pain management. Endometriosis is often considered to be an “estrogen-dependent” disease, but continued research aligning sex hormone with disease pathophysiology and patient outcomes is required to better interpret this complex relationship. Local tissue mechanisms more clearly implicate estrogen as a key factor in endometrial lesion growth and activity (*20–22*). Ectopic lesions aberrantly produce estrogen and have an increased abundance of *ESR2* relative to tissue from eutopic endometrium (the inner lining of the uterus), which lesions resemble. *ESR1* is less abundant in endometrial lesions relative to eutopic endometrium (*20, 23–27*). On the other hand, the role of systemic serum estrogen in endometriosis is less certain; systemic estradiol in non-medicated endometriosis patients was not different relative to healthy individuals (*28*), and another study identified that most serum estrogens and their derivatives are not associated with endometriosis pain aside from one particular derivative (*29*).

Regardless of whether local or systemic estrogen is more mechanistically implicated, it remains uncertain how much sex hormones in general can significantly impact clinical endometriosis pain outcomes. Additionally, direct assessments of sex hormone factors on the tissue on which lesions implant, most commonly the peritoneum (the serous membrane of the abdomen and pelvis), has not been deeply explored. Lesion-free peritoneum may itself contribute to the pain phenotype, warranting an independent evaluation and comparison to anatomically similar lesions. Ultimately, identifiable patterns of estrogen receptor levels across endometrial lesions, lesion-free peritoneum, as well as serum estrogen levels could prove to be critical determinants of endometriosis pain.

Here, we examined estrogen signaling in relation to clinical pain across these local (lesion and peritoneum) and systemic (serum) compartments. We aimed to 1) characterize estrogen and its receptors in patient-paired tissue biopsies and 2) examine whether clinical pain outcomes are influenced by systemic estrogen levels and/or estrogen receptor expression in lesions and patient-paired peritoneum tissues. Additionally, we performed exploratory analyses on other sex hormones (progesterone, prolactin, and testosterone) possibly involved in the disease collected via serum and their corresponding receptor expression levels in lesions and peritoneum (progesterone receptor, *PGR*; androgen receptor, *AR*; and long- and short-form prolactin receptor, *PRLR*-L and *PRLR*-S, respectively) (*15, 28, 30–37*). Together, these provide a map of the endocrine microenvironment in the context of pain associated with endometriosis.

## RESULTS

### Endometriosis lesion and peritoneum tissues display distinct sex hormone factor abundance

In a sample of 38 participants who underwent endometriosis resection surgery (see participant characteristics in **Table 1**), biopsies were collected from endometriosis lesions and lesion-free peritoneum as a control. To assess whether the local hormonal environment differs by disease status, patient-paired peritoneal and lesion tissues were analyzed for hormone receptor expression by real-time quantitative PCR (qPCR). Paired samples *t*-tests were used to compare differences in receptor levels between tissues. In addition, Pearson’s correlations were used to examine the within-subjects relationship between hormone receptor levels in peritoneum and lesions. Since endometriosis is hypothesized to be an estrogen-dependent disease, estrogen-related measures were chosen *a priori* as the main independent variables.

**Table 1.** Participant characteristics.

| **Variable** | **Total**<br>**participants**<br>**(*n* = 60)** | **Biopsied**<br>**participants**<br>**(*n* = 38)** | **Non-biopsied**<br>**participants**<br>**(*n* = 22)** |
| --- | --- | --- | --- |
| Age (years) | 31.78 (1.02) | 29.87 (1.19) | 35.09 (1.72) |
| BMI | 30.72 (1.38) | 28.61 (0.92) | 34.53 (3.39) |
| Race (white) | 53 | 32 | 21 |
| Ethnicity (Non-Hispanic) | 54 | 32 | 22 |
| Proper menstruation | 31 | 20 | 11 |
| Luteal | 13 | 8 | 5 |
| Follicular | 7 | 4 | 3 |
| Actively Menstruating | 11 | 8 | 3 |
| Altered menstruation | 29 | 18 | 11 |
| Phase impacted by hormone<br>medication usage | 14 | 10 | 4 |
| Other | 15 | 8 | 7 |
| Pain Medication Usage | 19 | 12 | 7 |

Variables are either reported as mean (standard error) or frequency. Proper menstruation indicates participants who were in the luteal, follicular, or actively menstruating phase of their menstrual cycle. Altered menstruation indicates participants who either had their cycles impacted by medication usage or “other” reasons such as irregular periods (>45 days since last period), surgical impact (e.g., hysterectomy or uterine ablation stopped menstruation), or unknown reasons.

The abundance of *ESR1* was significantly higher in lesions compared to the peritoneum (**Fig. 1A** and **Table 2**). Interestingly, there was a strong positive correlation between *ESR1* level in lesions and peritoneum, indicating that greater *ESR1* density in peritoneum is related to greater transcript levels in the lesion (**Fig. 1A′**). Conversely, *ESR2* expression did not differ by tissue source (**Fig. 1B**), although the expression within peritoneum and lesion tissue was significantly correlated (**Fig. 1B′**). Of note, *ESR1* was more highly expressed than *ESR2* in both lesions and peritoneum (**Supp. Fig. 1**). These data demonstrate distinct estrogen receptor-subtype differences in transcript level between lesion-free peritoneum and lesions. Notably, the large variability of expression values across individuals (**Fig. 1A′, 1B’**) also highlights substantial between-subjects variability in basal transcript levels.

**Fig. 1.**
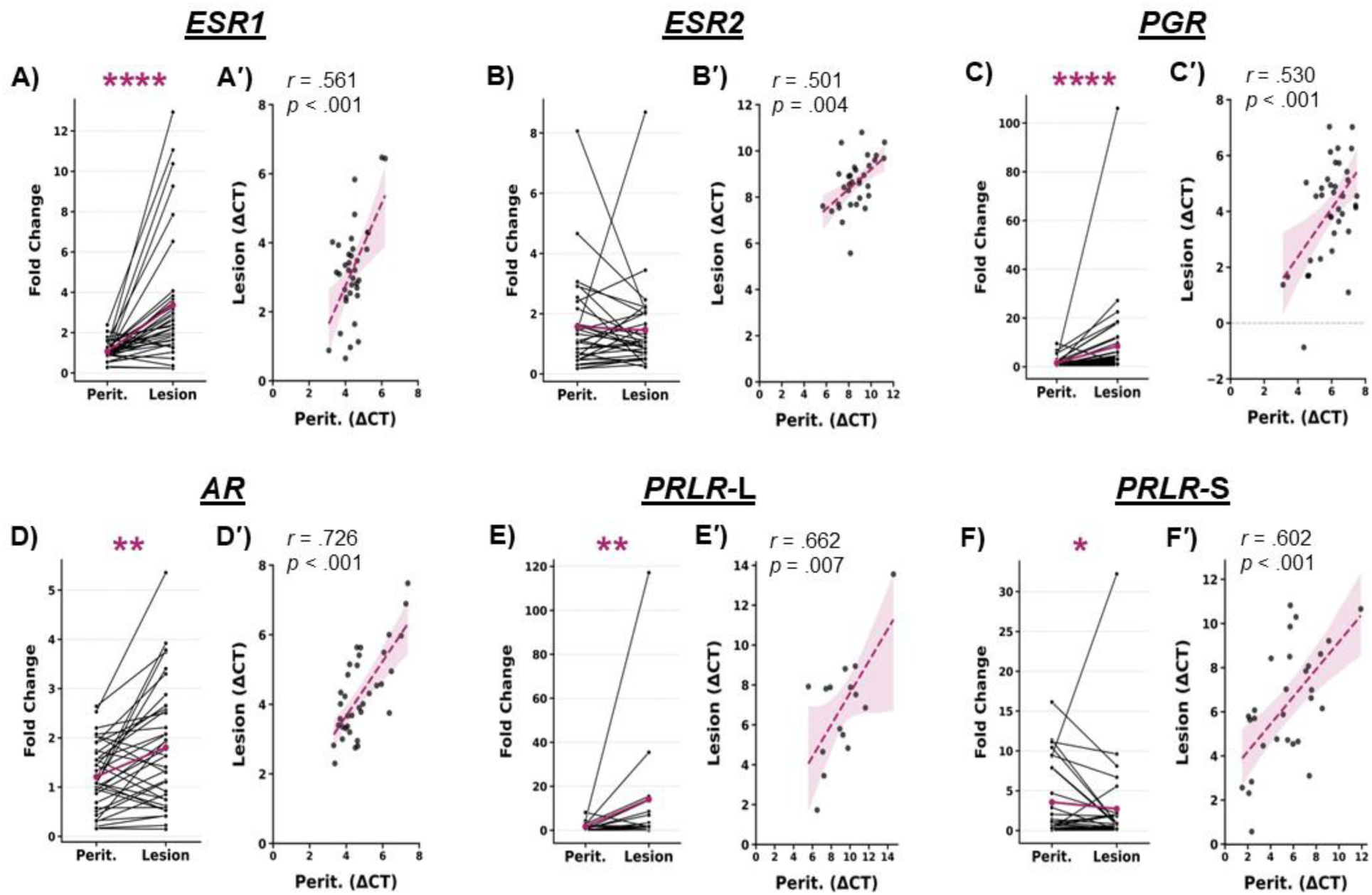
Analysis of local sex hormone levels and relationships between peritoneum and lesion of the same person. Spaghetti plots for B-G show the fold change relative to the mean of control values for both peritoneum and lesion expression. Correlation plots use ΔCT to show the relationship between expression levels in lesions and peritoneum. **A)** *ESR1* is enriched in lesions, and **A′)** levels are associated between tissues. **B)** There is no difference in *ESR2* between peritoneum and lesions, yet **B′)** there is a significant correlation between levels in each tissue. **C)** There are higher levels of *PGR* expression in lesions, with **C′)** positive association between each tissue type. **D)** *AR* is enriched in lesions, and **D′)** levels are associated between tissues. **E)** *PRLR-*L is higher in lesions, and **E′)** the levels between tissues are positively correlated. **F)** *PRLR-*S is lower in lesions compared to peritoneum, yet **F′)** the between tissue association remains. \**p* < 0.05, \*\**p* < 0.01, \*\*\**p* < 0.001. *Perit = peritoneum*.

**Table 2.** Paired samples *t*-tests comparing local hormone levels between healthy peritoneum and endometrial lesion tissue.

| Variable | *n* | Tissue | Mean | SEM | *t* | *p* | Cohen's *d* |
| --- | --- | --- | --- | --- | --- | --- | --- |
| *ESR1* | 36 | Lesion | 3.20 | 0.54 | -6.16 | < .001 | -1.03 |
|  |  | Peritoneum | 4.38 | 0.811 |  |  |  |
| *ESR2* | 32 | Lesion | 8.60 | 0.20 | -0.03 | .976 | -.005 |
|  |  | Peritoneum | 8.61 | 0.23 |  |  |  |
| *PGR* | 37 | Lesion | 4.06 | 0.29 | -7.60 | < .001 | -1.25 |
|  |  | Peritoneum | 5.95 | 0.18 |  |  |  |
| *AR* | 36 | Lesion | 4.28 | 0.20 | -3.46 | .001 | -.577 |
|  |  | Peritoneum | 4.79 | 0.19 |  |  |  |
| *PRLR-L* | 15 | Lesion | 6.88 | 0.71 | -4.07 | .001 | -1.05 |
|  |  | Peritoneum | 9.11 | 0.59 |  |  |  |
| *PRLR-S* | 29 | Lesion | 6.30 | 0.49 | 2.09 | .046 | .388 |
|  |  | Peritoneum | 5.39 | 0.48 |  |  |  |

*Note.* qPCR data is presented as the mean ΔCT value for each gene where a lower number represents higher expression. *ESR1* = *estrogen receptor alpha*. *ESR2* = *estrogen receptor beta*. *PGR* = *progesterone receptor*. *AR* = *androgen receptor*. *PRLR-L* = *long prolactin receptor isoform*. *PRLR-S* = *short prolactin receptor isoform*.

Since endometriosis can be modulated not only by estrogen but several sex hormones (*28*), the expressions of *PGR*, *AR*, *PRLR*-L, and *PRLR*-S were also evaluated as exploratory variables (**Table 2**). The expressions of *PGR*, *AR*, and *PRLR*-L were significantly higher in lesions compared to peritoneum (**Fig. 1C-E**), whereas *PRLR*-S was significantly lower in lesions compared to peritoneum (**Fig. 1F**). *PRLR*-S was more abundant and more frequently detected by qPCR than *PRLR*-L in peritoneum, while no isoform difference was observed in lesion tissue (**Supp. Fig. 1**). All receptors showed significant correlations between their expression levels in the lesion and peritoneum (**Fig. 1C′-F′**).

Analyses between factors in peritoneum or lesion revealed interesting patterns of expression. We found strong associations between *ESR1*, *PGR*, *AR,* and *PRLR*-L levels in lesions but fewer associations between factors in peritoneum were identified (**Fig. 2**). These results imply a coordinated sex hormone receptor response which is enriched in lesion tissue relative to peritoneum in endometriosis. Given the differences in sex hormone receptor densities between tissues, and the potential that non-lesion peritoneum tissue might contribute to clinical pain, we accounted for both the abundance of receptors in each tissue as well as the change in receptor abundance between tested tissue within each participant in subsequent analyses.

**Fig. 2.**
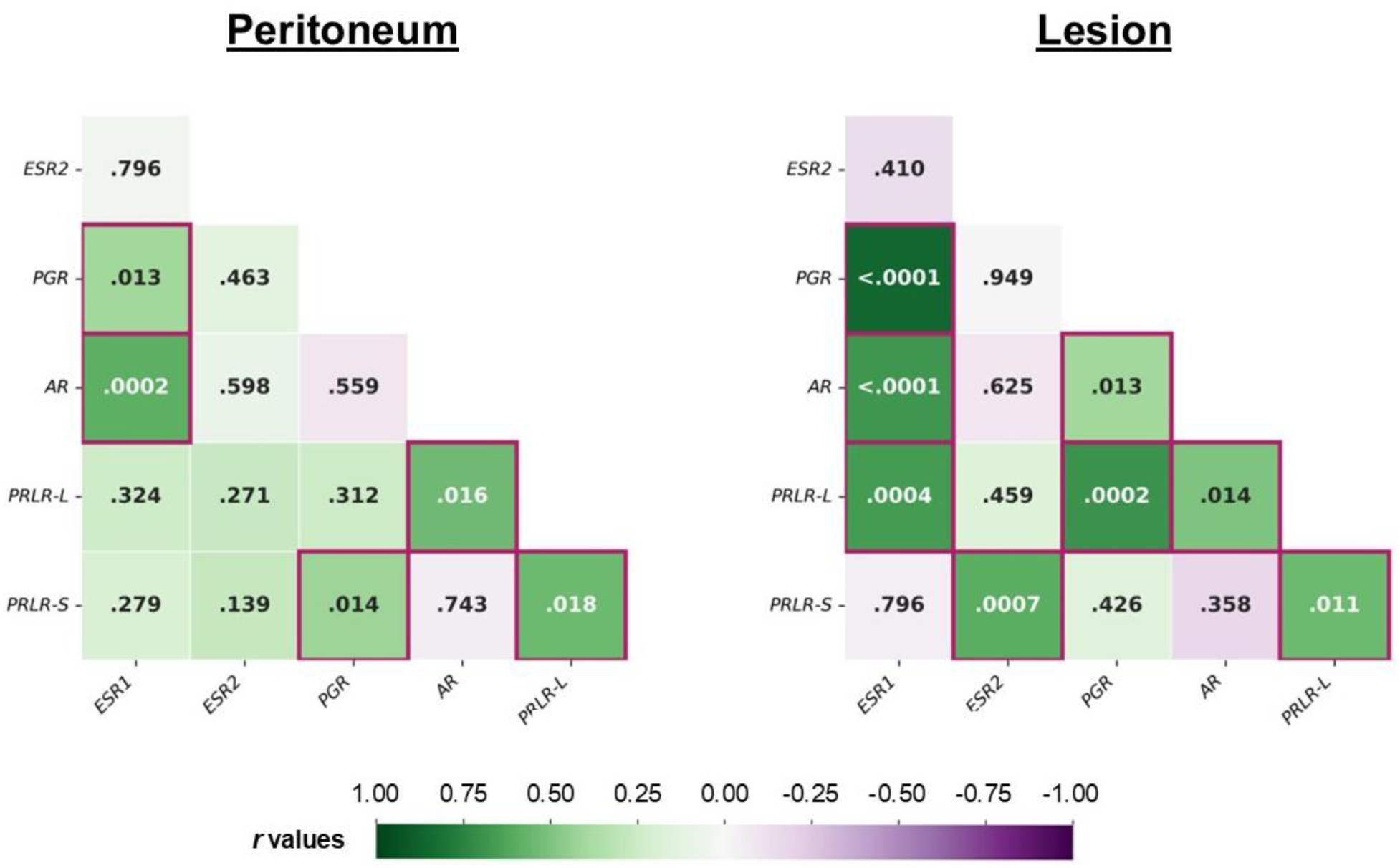
Correlations between receptor expression levels in peritoneum (left) and lesion (right) tissues. Shade of the box indicates the *r*-value. Number within the box is the *p*-value. Boxes are outlined to highlight significant associations. *ESR1 = estrogen receptor alpha. ESR2 = estrogen receptor beta. PGR = progesterone receptor. AR = androgen receptor. PRLR-L = long prolactin receptor isoform. PRLR-S = short prolactin receptor isoform*.

### Tissue hormonal factors are poor predictors of endometriosis pain

Before the surgery, participants completed a study visit which included questionnaires assessing sociodemographic and clinical pain-related variables. The Endometriosis Health Profile-30 (EHP-30) assesses the overall impact of endometriosis on different aspects of life, but specifically, the “pain” subscale focusing on the impact of pain related to endometriosis was used in the present analyses. The Patient-Reported Outcomes Measurement Information System (PROMIS) Pain Interference scale was used to broadly examine how pain impacts daily functioning. Lastly, the level of endometriosis/pelvic pain intensity in the last 30 days was rated from *0 = no pain* to *10 = worst pain imaginable*.

To directly compare between tissue types, the transcription fold change in lesions relative to same participant peritoneum (2^-ΔΔCT) was calculated. Linear regressions were run to examine whether the factor change between tissues is related to clinical pain while controlling for menses phase classification, age, and pain medication usage. In addition, linear regression of lesion or peritoneum factor levels (ΔCT) as separate predictors were run to determine unique relationships with clinical pain while controlling for the same covariates. Estrogen-related factors were chosen *a priori* as the main predictors of interest while the other receptors were considered exploratory. No significant relationships were found between any of the clinical pain measures and the relative change in sex hormone receptors (*ESR1*, *ESR2*, *AR*, *PGR*, *PRLR*-L, *PRLR*-S; **Fig. 3A, Suppl Table 1**). Almost all the regressions using receptor expression in lesions or peritoneum as separate predictors failed to reach significance except for *ESR1* and *PGR* (**Fig. 3B, Suppl Table 1**). These factors were found to be negatively associated with pain intensity in the last 30 days, where greater pain intensity was predicted by lower lesion expression of *ESR1* (*ΔR^2^* = .138, *ΔF*_1,27_ = 4.46, *p* = .044, *β* = .403, 95% CI = [.095, .944]) (**Suppl Fig. 2A**) and *PGR* (*ΔR^2^*= .261, *ΔF*_1,27_ = 9.91, *p* = .004, *β* = .562, 95% CI = [.144, .897]) (**Suppl Fig. 2B**). Because estrogen was selected a priori as the primary independent variable, we applied a correction for multiple comparisons. Following correction, the association between pain intensity and ESR1 expression in endometriosis lesions was no longer statistically significant.

**Fig. 3.**
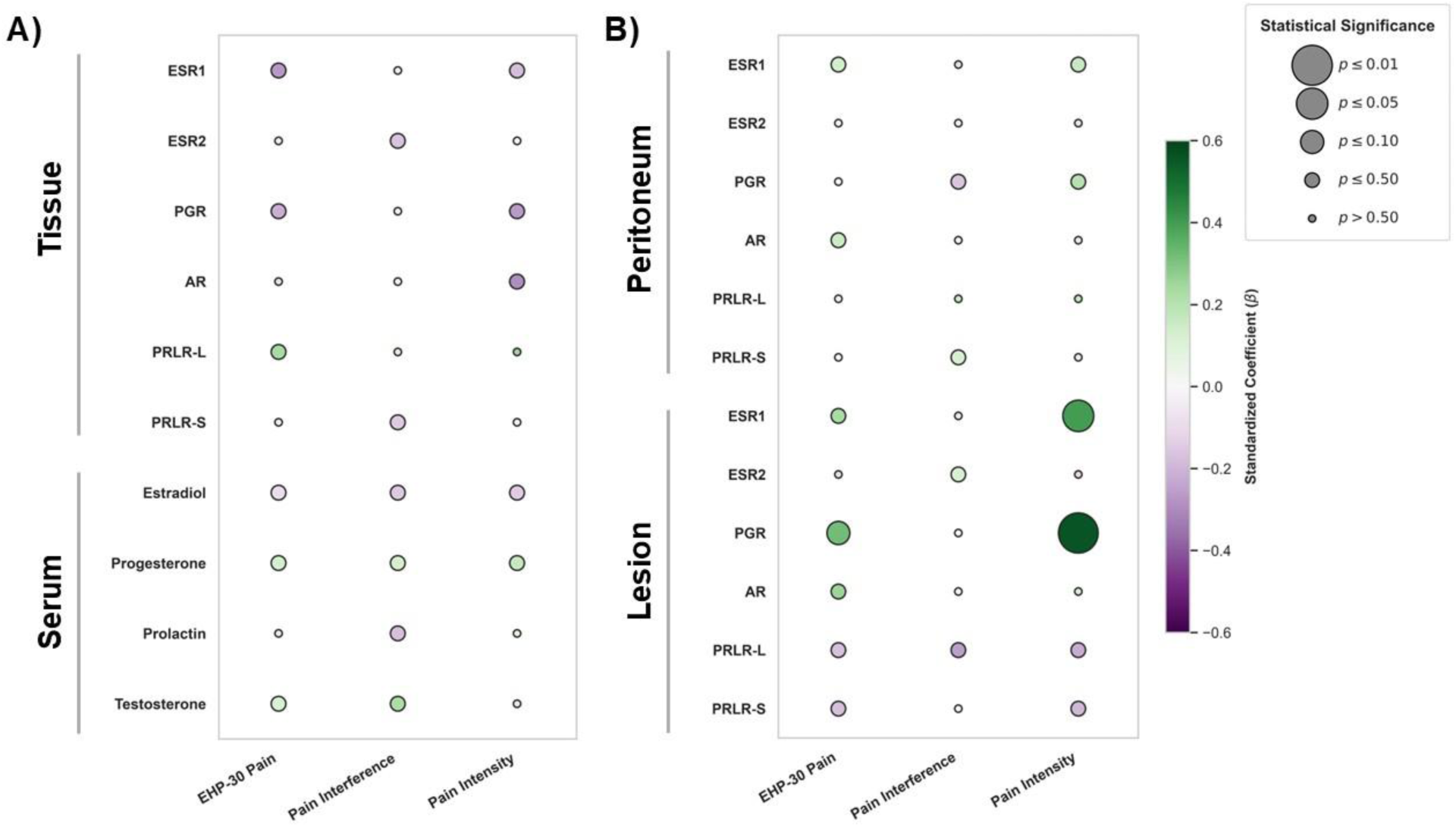
Linear regressions with sex hormone levels predicting clinical pain outcomes. Size of the dots indicates the statistical significance, representing the predictor’s significance beyond the influence of covariates. Color tone of the dots indicates the standardized coefficient (β) values, representing the predictor’s contribution to the outcome beyond the influence of covariates. The outcome of EHP-30 pain represents the pain subscale of the Endometriosis Health Profile 30 questionnaire. The outcome of pain interference is the PROMIS Pain Interference t-score. The outcome of pain intensity is the rating of average pelvic pain intensity in the last 30 days on a visual analogue scale (VAS) where *0 = no pain* to *10 = worst pain imaginable*. Covariates included age, menstrual cycle phase classification, and pain medication use. Regressions using serum sex hormones as predictors included the time of blood draw as an additional covariate. A) Predictors include the transcription fold change of sex hormone receptors (from *ESR1* to *PRLR-*S) and serum sex hormones (from estradiol to testosterone). None of these factors significantly predicted any clinical pain outcomes. B) Predictors include the tissue level of each receptor in peritoneum (top half) and lesions (bottom half). Both *ESR1* and *PGR* expression in endometrial lesions significantly predicted pain intensity, indicating that less receptors present in lesion tissue may contribute to greater endometriosis/pelvic pain intensity. However, significance of *ESR1* did not survive FDR-correction (*p* = .308). No other factors significantly predicted any clinical pain outcomes. *ESR1 = estrogen receptor alpha. ESR2 = estrogen receptor beta. PGR = progesterone receptor. AR = androgen receptor. PRLR-L = long prolactin receptor isoform. PRLR-S = short prolactin receptor isoform*.

These data support that expression of *ESR1* and *PGR* specifically in lesions may have an influence on endometriosis/pelvic pain intensity since neither the peritoneum expression nor the relative change in receptor abundance between lesion and peritoneum were significant predictors. It is possible that pathological features of lesions alter *ESR1* and *PGR* expression (or cell populations expressing them) which can modulate clinical pain outcomes.

### No relationships between serum sex hormone levels and endometriosis pain were identified

An additional 22 women with preexisting endometriosis diagnosis who did not undergo surgery during the study were included for a total sample size of 60 participants (see participant characteristics in **Table 1**). During the study visit or prior to endometriosis lesion resection surgery for some participants, venous blood was collected for serum hormone quantification. To determine whether serum hormone levels are related to clinical pain, separate linear regression models were conducted controlling for menses phase classification, age, pain medication usage, and the time of blood draw. Serum estradiol was chosen *a priori* as the main independent variable while the other systemic hormones (prolactin, progesterone, and testosterone) were considered exploratory variables. No relationships were found between any of the serum sex hormone levels and any of the clinical pain outcomes (**Fig. 3A, Suppl Table 1**).

## DISCUSSION

This study uniquely associated patient-paired tissue transcript levels of sex hormone receptors and systemic sex hormones with clinical pain outcomes in individuals with endometriosis. To our surprise, no serum sex hormone levels nor relative receptor expression levels in lesions compared to peritoneum were related to any of the clinical pain outcomes. However, the transcript levels of *ESR1* as well as *PGR* in lesions were negatively related to endometriosis/pelvic pain intensity, suggesting that greater pain was associated with lower expression levels of *ESR1* and *PGR* in endometrial lesions, although after false discovery rate (FDR) correction the association with *ESR1* was no longer significant. Taken together, these data support that sex hormone factors may relate to endometriosis pathophysiology, but their relationships with clinical pain outcomes are much more complex being both tissue- and factor-specific.

### Molecular distinctions between non-lesion peritoneum and endometriosis lesions

It is well established that ectopic endometrial lesions contain distinct cell types relative to other anatomical sites such as the eutopic endometrium (*24–26*). Although lesions are the primary focus of research studies due to their pathology, molecular investigations have also revealed distinctions in eutopic endometrium and non-lesion peritoneum between endometriosis patients and healthy controls (*24–27, 38*). We found a striking relationship supporting that matched peritoneum and lesion tissue are closely related. Lesions showed a higher abundance of *ESR1* and other sex hormone receptors than peritoneum, with no significant difference in *ESR2* between tissue types, yet receptor levels in the two tissues were significantly correlated within participants. Given that prior studies have reported a significant difference in peritoneum in addition to lesions between patients and controls (*38*), these results further suggest that ‘healthy’ peritoneum may itself be affected by the disease. This could influence disease pathophysiology but not pain as the factors we tested in peritoneum did not relate to clinical pain outcomes at the transcriptional level.

In the context of these findings, it is noteworthy that *ESR2* levels are not different between tissue types. This observation diverges from classic reports which found a profound increase in expression of stromal *ESR2* when comparing eutopic endometrium to ectopic lesions (*39*). Indeed, single-cell sequencing data support these reports where *ESR2* expression is virtually absent in eutopic endometrium (*27*). Therefore, the lack of differential *ESR2* signals in our dataset is likely due to the presence of *ESR2* expression in peritoneum. Collectively, these data support a multimodal hypothesis that describes contributions of both the origin of endometrial-like cells (*4, 13*) along with implantation site tissue to promote lesions and disease (*38*). Utilizing bioinformatics and transcriptomics to integrate the signatures of cellular origin with implantation site tissue may therefore offer a powerful approach to better define the specific cellular subsets and gene expression that are unique to lesions.

### Associations across sex hormones tested

Interestingly, we found that *ESR1* and *AR* expression is significantly correlated within peritoneum, but lesions exhibit stronger relationships between multiple sex hormone receptors including *ESR1, PGR,* and *AR.* These function as classical nuclear receptors and therefore share some downstream transcriptional and regulatory mechanisms (*40–42*). Notably, recent evidence has identified that the ratio of these three transcripts is predictive of tumor subtypes and patient outcomes in breast cancer, another sex-hormone-dependent disease (*42*). Previous single-cell RNA sequencing studies comparing eutopic endometrium to ectopic lesions reveal lesion enrichment of *AR* and *PGR* in stromal fibroblasts and *ESR1* in epithelium. Furthermore, in the eutopic endometrium, there is robust expression of *ESR1*, *AR*, and *PGR* in the same compartment of stromal cells (*27*). It is plausible that the heightened association found between these genes within lesions is driven by endometrial-like stromal cells found within lesions. Taken together, it is clear there is an important role of other sex hormones, specifically progesterone and androgens, in endometriosis pathophysiology beyond the influence of estrogen. Consequently, further examining the specific cell types expressing these factors and elucidating how they differ in non-lesion peritoneum and lesion tissues is of critical importance in understanding endometriosis.

### Pain as a determining factor

Pain is a complex sensation involving a multitude of processes in both peripheral and central nervous systems, including sex hormones as important modulators of pain sensation. Of interest, estrogen directly modulates nociception at primary sensory neurons with extensive expression of ERα and ERβ on sensor neurons in the dorsal root ganglion (*43*). Evidence for the role of estrogen in inflammation and nociception is mixed and context dependent (*38, 44*), although high concentrations of estradiol activate inhibitory pathways in the spinal cord to engage antinociceptive pathways, while low concentrations exhibit pronociceptive properties (*45, 46*). Indeed, since estrogen and other sex hormone receptors are located in various brain areas that are involved in pain processing, these hormones can have a diverse impact on pain (*47–49*).

To provide a more detailed understanding of how peripheral sex hormones can impact endometriosis pain, we examined whether tissue or serum hormones are related to endometriosis-related pain impact (EHP-30 pain subscale), endometriosis/pelvic pain intensity, and pain interference. Surprisingly, almost no relationships with sex hormone levels or receptor levels were found with any of the pain measures, suggesting that other factors may have a greater impact on the severity of pain. However, we did detect an association with pain intensity and lesion expression, specifically of *ESR1* and *PGR although the relationship with ESR1 was no longer significant after FDR correction for multiple comparisons*. Interestingly, lower abundance of both transcripts was related to greater endometriosis/pelvic pain intensity in the last 30 days. Further studies replicating the results and mechanistic studies to determine the causality of these associations are necessary. Still, these data align with previous clinical observations. Estrogen and progestins are commonly prescribed to manage endometriosis pain (*15, 17, 18*), yet progesterone resistance commonly occurs (*33*). In these patients, progestins lose effectiveness in reducing pain, possibly due to a reduction in *PGR* abundance in tissue (*33*). Importantly, in our analyses, we controlled for participant hormonal medication usage. As discussed above, *ESR1* and *PGR* are highly enriched (**Fig. 1**) and uniquely associated with one another in lesions compared to control tissue (**Fig. 2**), suggesting a coordinated response between the factors which might impact clinical pain. Mechanistic or interventional studies to increase *ESR1* and *PGR* in lesions would determine whether these factors contribute to pain by regulation of endometrial-like cells, inflammatory cells, or innervating nerve fibers (*50, 51*).

In contrast to these localized tissue dynamics, systemic levels of estradiol and other sex hormones demonstrated no relationship to clinical pain outcomes. This disconnect may be due to fluctuations characteristic of circulating serum hormones relative to the more stable profiles found in tissue (*52*). Indeed, endometriosis itself contributes to systemic sex hormone alterations, mostly irrespective of individual pain thresholds (*28*). It may also be that absolute hormone levels taken at a single timepoint in the disease are insufficient to explain the complex pain phenotypes. Instead, longitudinal evaluations tracked across disease progression and treatment may be required to uncover latent relationships between systemic sex hormones and pain reports.

Of similar clinical interest are the factors that did not relate to reported pain. *PRLR*-L is lowly expressed overall and is found significantly lower than *PRLR*-S in peritoneum (**Supp. Fig. 1**). While *PRLR*-S is enriched in peritoneum (**Fig. 1**), its levels and directions are highly variable between participants, an interesting result since *PRLR*-S on female sensory neurons increases neuronal excitability (*53–55*). Furthermore, *ESR2* is unchanged between tissue types (**Fig. 1**), unlike *ESR1*, but has been associated with pain/nociception (*56–58*). Both *PRLR*-S and *ESR2* display large variability in our dataset, which may relate to other aspects of disease pathophysiology given the lack of significant relationships with clinical pain.

### Limitations and strengths

There are several limitations to this study. The regression analyses of the non-estrogen sex hormone factors were exploratory and include no correction for multiple comparisons, so the results of these regression models should be treated with caution and need to be validated in future studies. Additionally, we relied on select mRNA transcripts which may not directly align to protein levels. Further, there are multiple instances in which there was no signal detected in qPCR analyses (**Supp. Fig. 1**). While these cannot be directly analyzed, it is worth noting that the incidence “not detected” was higher in non-lesion peritoneum compared to lesions, suggesting that the actual relative differences between tissues for some genes may be greater than reported.

The strengths of the study include comparisons of multiple key sex hormones in the same participants across three different sample sources with the ability to relate these tissue and serum levels to clinical pain outcomes. Since tissue samples were collected at the same time for each participant, variation due to ovarian cycle or medications were controlled by within-participant testing. The inclusion of non-lesion peritoneum provides an important advancement as a tissue of interest and acts as an internal reference revealing both between-tissue and between-participant differences in the factors tested. Together, these data evaluate both common implantation sites and patient-paired lesions to determine sex hormone signaling in endometriosis. Overall, we found sex hormone factor tissue and serum levels are overall poor predictors of endometriosis-associated pain and disability.

### Conclusions

In this study, we examined relationships and differences between key sex hormone factors within grossly normal peritoneum where lesions could form and existing endometrial lesions, finding distinct endocrine signatures between the two. We uncovered significantly greater abundance of *ESR1* and multiple sex hormone receptors in lesions compared to same participant peritoneum, with a notable lack of difference in *ESR2* between tissues. Overall, we found sex hormone tissue and serum levels are poor predictors of endometriosis-associated pain and disability. Additional studies are necessary to validate these findings. Collectively, our findings provide evidence of distinct, site-specific molecular programming at both the host peritoneum and ectopic lesion which may dictate aspects of disease pathophysiology. Additional future research is vital to improve our understanding of the factors driving worse pain outcomes for patients and to develop more direct treatments for those most vulnerable to painful endometriosis.

## MATERIALS AND METHODS

Washington University School of Medicine’s IRB committee approved all the procedures, and all participants signed informed consent. In case of participants under 18, both legal guardians signed the consent, and the participant signed the assent. Preregistration was completed on ClinicalTrials.gov (NCT06101303).

### Participant Recruitment and Characteristics

Patients scheduled for laparoscopic surgery for the diagnosis and excision of endometriosis in the Department of Obstetrics and Gynecology at Washington University School of Medicine were contacted. Inclusion criteria included individuals aged 12-45 years old scheduled for laparoscopic surgery for suspected or known endometriosis. Exclusion criteria included current pregnancy, lactation, or cancer diagnosis. Patients with other chronic pain conditions, psychiatric disorders, or those taking pain or hormonal medications were not excluded, as these factors are common in individuals with endometriosis and excluding them would limit the clinical relevance and generalizability of the findings. In addition, participants of ages 12-56 with a pre-diagnosis of endometriosis and not currently treated for diagnosed psychiatric, neurological, endocrinological, or developmental conditions were recruited using flyers around Washington University School of Medicine and other public locations around Saint Louis, Missouri, as well as via email using Washington University’s research volunteer database (see participant characteristics in **Table 1**).

### Study design

Participants completed a study visit (*n* = 60) which included questionnaires assessing sociodemographic and clinical pain-related variables. The study visit was before the surgery for participants undergoing surgery (*n* = 38). At the end of the study visit, a blood sample for the analyses of systemic sex hormone levels was collected. For some surgical participants, the blood sample for systemic hormone levels was collected prior to the initial incision but after undergoing general anesthesia.

When possible, biopsies of the endometriosis lesion and the lesion-free peritoneum were collected from participants during surgery. Surgeons identified endometriosis lesions which were histologically confirmed by pathology when possible. The anatomic location and macroscopic characteristics of the lesions were noted for evaluation of endometriosis-related pain.

### Pain-related measures

#### Endometriosis Health Profile-30 (EHP-30) (*59*)

This 30-question survey assesses self-reported quality of life in the last month. Answers range from *0 = never* to *4 = always*. The average of the questions was calculated, with higher scores indicating a worse quality of life due to endometriosis-related symptoms.

#### PROMIS Pain Interference (*60, 61*)

The PROMIS pain interference scale was used to determine how pain in the last week impacted their daily functioning. Questions are answered from *1 = never* to *5 = always* which is then summed and converted to a standardized T-score. Higher scores indicate a greater degree of interference.

### Pain intensity

Participants were asked to rate the level of their endometriosis/pelvic pain intensity in the last 30 days on a numeric rating scale from *0 = no pain* to *10 = worst pain imaginable*.

### Serum sex hormone level quantification

10 mL intravenous blood was drawn and immediately placed into serum separation tubes. Following blood collection, the tube was inverted and kept upright at room temperature for at least 30 minutes. After clotting, blood was centrifuged at 3000 rpm for 10 minutes within 60 minutes from collection. The serum was removed and stored at -20°C until transported to LabCorp (Laboratory Corporation of America® Holdings, USA) for processing. Serum levels for testosterone (ng/dL) and progesterone (ng/dL) were quantified by high-pressure liquid chromatography-tandem mass spectrometry (LC-MS/MS). Estradiol (pg/mL) was quantified by two-dimensional high-performance LC-MS/MS. Prolactin (ng/mL) was quantified by electrochemiluminescence immunoassay (ECLIA). For values lower than the quantifiable range, the minimum detection value was recorded.

### Biopsies collection

A biopsy of ‘healthy’ peritoneum on which no visible disease was noted was collected by the gynecologic surgeon for the study team. Lesions were then excised, ensuring that at least one specimen was sent to pathology. Biopsy characteristics were described by the surgeon prior to lesion resection. The tissues were immediately snap frozen on dry ice, and the biopsies were stored at -80°C until use.

### Tissue level quantifications

Tissue samples (5-30 mg) were processed for total RNA isolation. RNA was isolated using RNAqueous Total RNA Isolation kit (Invitrogen AM1912) according to manufacturer instructions. RNA quality and quantity were assessed by Nanodrop prior to cDNA generation using a High-Capacity cDNA Reverse Transcription Kit (Applied Biosciences 4368814) in 50 ng reactions. Real-time quantitative PCR (RT-qPCR) was performed in duplicate using Sybr Green (Biorad 1725271) and primers previously reported (*57*) for *ESR1* (Fwd: TGGGCTTACTGACCAACCTG, Rev: CCTGATCATGGAGGGTCAAA), *ESR2* (Fwd: AGAGTCCCTGGTGTGAAGCAA, Rev: GACAGCGCAGAAGTGAGCATC), *AR* (Fwd: TTGGATGGCTCCAAATCAC, Rev: GCAATGATACGATCGAGTTCC), *PGR* targets both isoforms A and B (Fwd: CGCGCTCTACCCTGCACTC, Rev: TGAATCCGGCCTCAGGTAGTT), and designed specifically for *PRLR*-L (Fwd: TCACTCCTTCTCCCTCTTTCT, Rev: GGGTCTGGCTGAACTATGTAAG; variant 1) and *PRLR*-S (Fwd: CATTGACCACAGTCCAGTAGAG, Rev: CCTTCACAAACACCTGCATAAC; variants 3, 4, 5). GAPDH (Fwd: ACCACCAACTGCTTAGCACC, Rev: CCATCCACAGTCTTCTGGGT) was used as an internal control. Technical duplicates were averaged.

Data is shown as either the delta cycle threshold (ΔCT) after normalization to GAPDH levels or fold change (2^-ΔΔCT) of lesion relative to either the mean of the non-lesion peritoneum (**Fig. 1**), or relative to same-participant non-lesion peritoneum (**Fig. 3**). -ΔCT is displayed for directional purposes in regressions (**Supp. Fig. 2**). A CT value equal to or greater than 34 was considered not detected (n.d.) (*62, 63*).

### Statistical Analysis

All statistical analyses were performed using SPSS (v. 31). Outliers more than 3 times the interquartile range were identified in all predictors and outcomes of interest and removed for each appropriate analysis involving the impacted variable. One participant exhibited outliers in *ESR1*, *PGR*, and *AR* fold change and was excluded from the corresponding analyses. For fold change, an additional two participants were excluded for *ESR1*, one for *PGR*, one for *ESR2*, and one for *PRLR*-S outliers. One participant was excluded for an outlier in peritoneal *ESR1* expression. For serum measures, three participants each were excluded from the *EST* and *PRL* analyses, and two from the *TTS* analyses. In total, 16 participants were excluded for the appropriate analyses. All outliers exhibited higher values than the rest of the sample. No outliers for clinical outcomes of interest or lesion receptor expression were identified. For participants with endometriosis lesions and control peritoneal tissue samples, paired-samples *t*-tests were used to compare tissue-specific differences in hormone receptor gene expression. Pearson’s correlation analyses were performed to assess within-subject associations between factor levels measured in lesion and peritoneal tissues.

The primary clinical pain outcome of interest was the degree of endometriosis-related pain as measured by the EHP-30 pain subscale. PROMIS pain interference and endometriosis/pelvic pain intensity in the last 30 days were included as clinical outcomes of interest. Separate linear regression models were used to determine whether the transcription fold change independently relate to clinical pain outcomes. Additional regression analyses were conducted using levels of each factor measured separately in lesion and peritoneal tissues to determine whether expression levels were associated with pain outcomes. Lastly, serum sex hormone levels were used as additional predictors of clinical pain in separate regression analyses. Given the predominantly exploratory nature of the linear regressions, multiple comparisons corrections were conducted for estrogen measures but not for the other hormones. To evaluate associations within each distinct clinical domain, Benjamini–Hochberg False Discovery Rate (FDR) corrections were conducted separately for each pain outcome (EHP-30 pain subscale, pain interference, and pain intensity).

For each outcome, *p*-values across all six corresponding models (*ESR1* and *ESR2* across lesion, peritoneum, and fold-change measures) were combined into a single family. Statistical significance was defined as an FDR-adjusted *q*-value < .05. Both uncorrected *p*-values and FDR-adjusted *q*-values are reported for all models.

Covariates for separate linear regressions included age, menstrual cycle phase classification, and pain medication use (*0 = not taking any, 1 = taking at least one pain medication*). For the menses phase classification, participants were classified by their menstrual phase during the study visit based on the first date of their last period, with those in the luteal phase, follicular phase, or active menstrual phase categorized into separate groups. Those taking any type of hormonal medication were classified into a single group given how hormonal alterations influence the menstrual cycle (*64, 65*). Participants actively menstruating were classified together regardless of whether hormonal medication is regularly taken. Participants with any other reason for altered menstrual cycling (e.g., hysterectomy, unknown reasons) were classified in a single group separate from the others. The time of blood draw recorded on the 24-hour clock and turned into a decimal value was included as a covariate for regressions using serum sex hormones as predictors.

Figures were created using Python 3 or GraphPad. Python code used to generate figures was aided with assistance from Google AI.

## List of Supplementary Materials

Fig. S1 to S2 for multiple supplementary figures

Table S1

## Acknowledgments

The authors would like to thank Joel Brown, Alana McMichael, Peyton Blunt, and Elizabeth Mulat for assistance in recruitment and data collection. Google AI was used for assistance with Python code to create some figures.

## Author contributions

Conceptualization: AJD, HNA, WTR, RWG

Methodology: AJD, HNA, WTR, RWG

Investigation: AJD, RLCO, JM, MD, WTR, EB, HRH

Formal analysis: RLCO, AJD, HNA

Visualization: RLCO, AJD

Funding acquisition: AJD, RLCO, HNA, WTR

Project administration: HNA

Supervision: HNA, RWG

Writing – original draft: AJD, RLCO, HNA

Writing – review & editing: JM, HRH, MD, EB, TJB, RWG, WTR, BPK

## Data and materials availability

Because data is still being collected and analyzed for additional study aims, the data supporting these findings will not be made available until analyses for all study aims are completed.

## Supplemental Materials

**Fig. S1.**
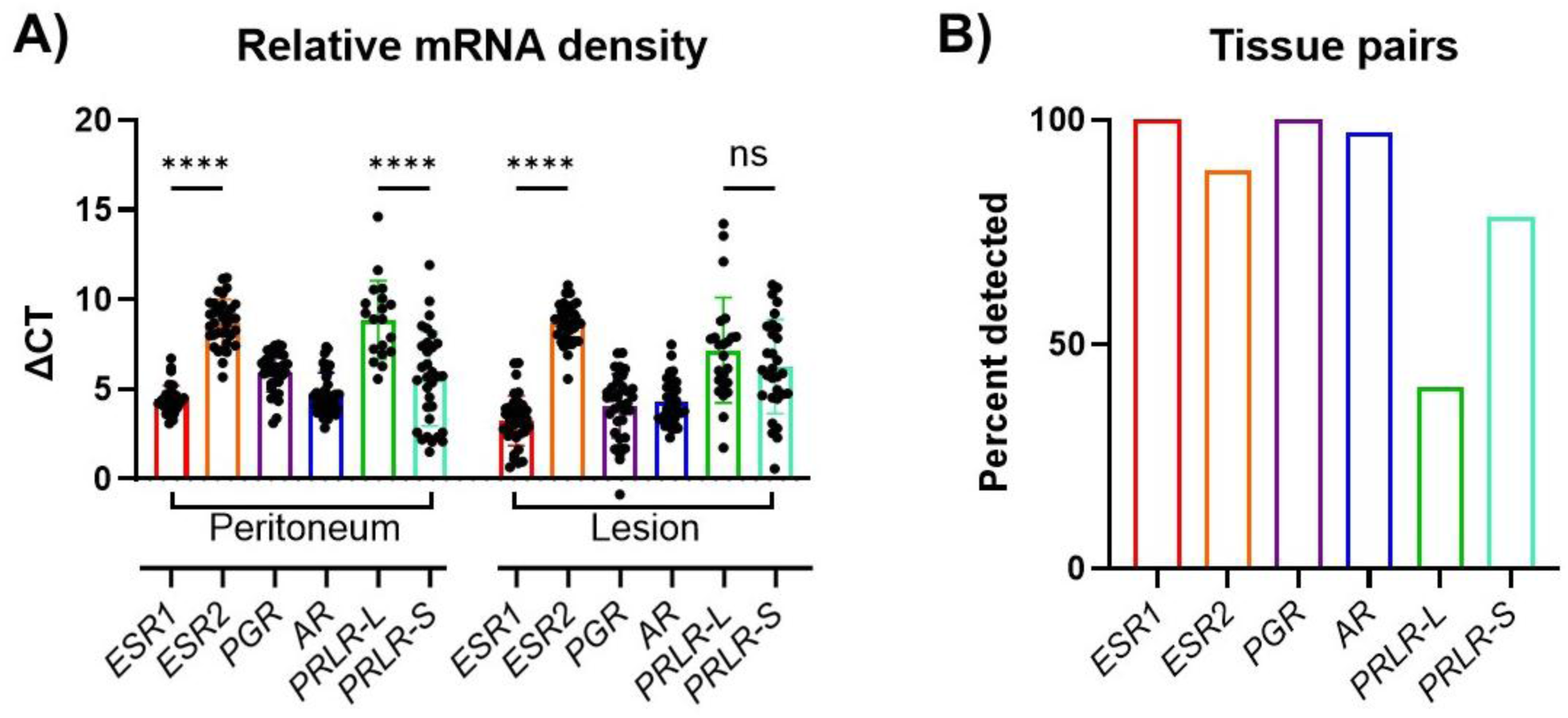
PCR detection metrics. **A)** Analyses between relevant factor expression level within peritoneum or lesion tissues. *ESR1* is more highly expressed than *ESR2* in both tissues, while *PRLR*-S is more highly expressed than *PRLR*-L only in peritoneum (\*\*\*\**p* < 0.0001, paired student *t*-test). **B)** The percentage of samples detected in both peritoneum and lesions tissues. Graphed data are ΔCT relative to GAPDH where a lower value indicates higher expression. Data shown as the mean±SEM.

**Fig. S2.**
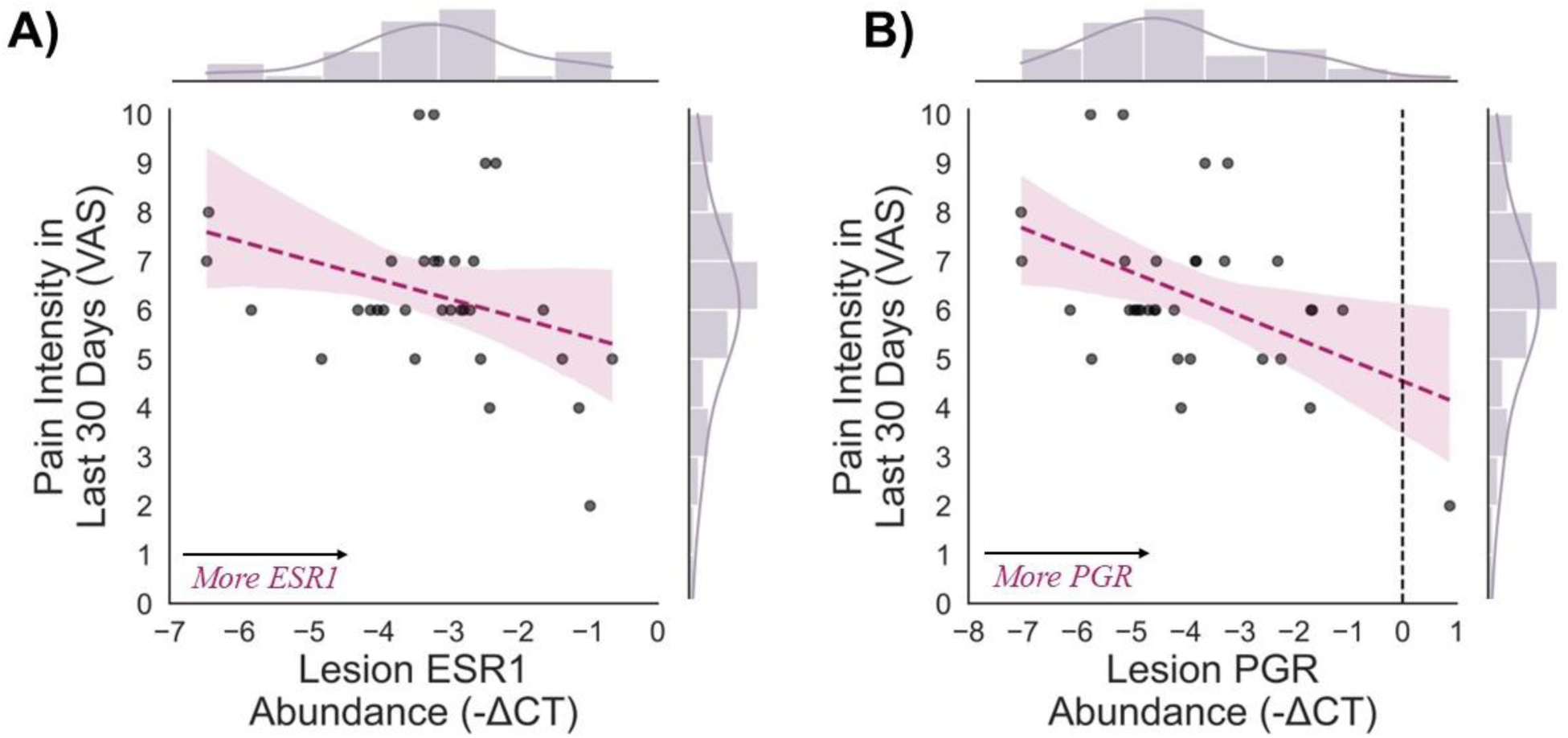
Linear regression examining the relationship between endometrial tissue expression and pain intensity. Lesion abundance is displayed as the negative ΔCT to better visually understand the relationship between the factors. **A)** Lower expression of *ESR1* receptors in lesion tissue significantly predicts worse pelvic pain intensity in the last 30 days beyond the influence of covariates (*ΔR^2^*= .138, *ΔF*_1,27_ = 4.46, *p* = .044, *β* = .403, 95% CI = [.095, .944]). However, this did not survive FDR-correction (*p* = .308). **B)** Lower expression of *PGR* receptors in lesion tissue significantly predicts worse pelvic pain intensity in the last 30 days beyond the influence of covariates (*ΔR^2^* = .285, *ΔF*_1,28_ = 11.48, *p* = .002, *β* = .583, 95% CI = [.185, .914]).

**Supplemental Table 1.**
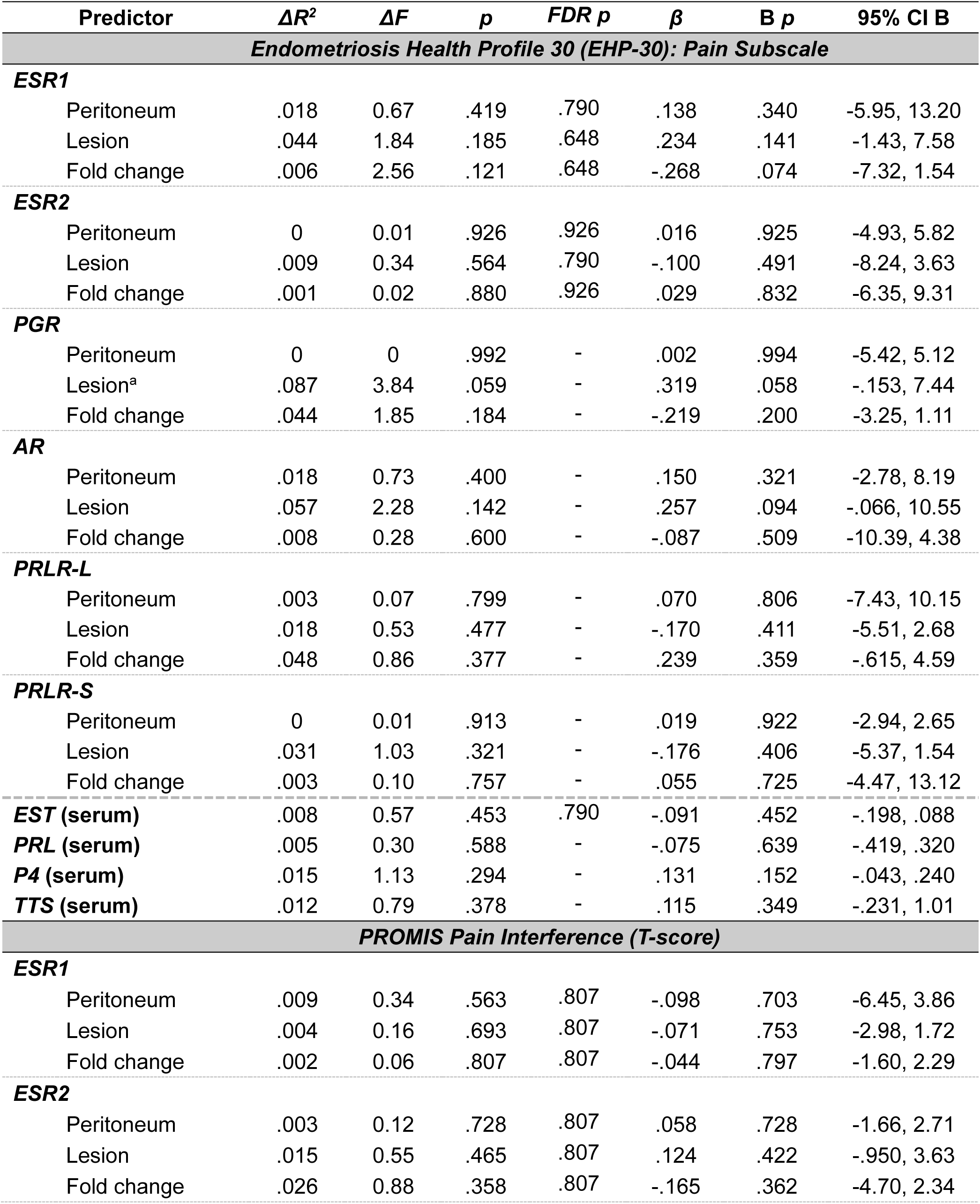
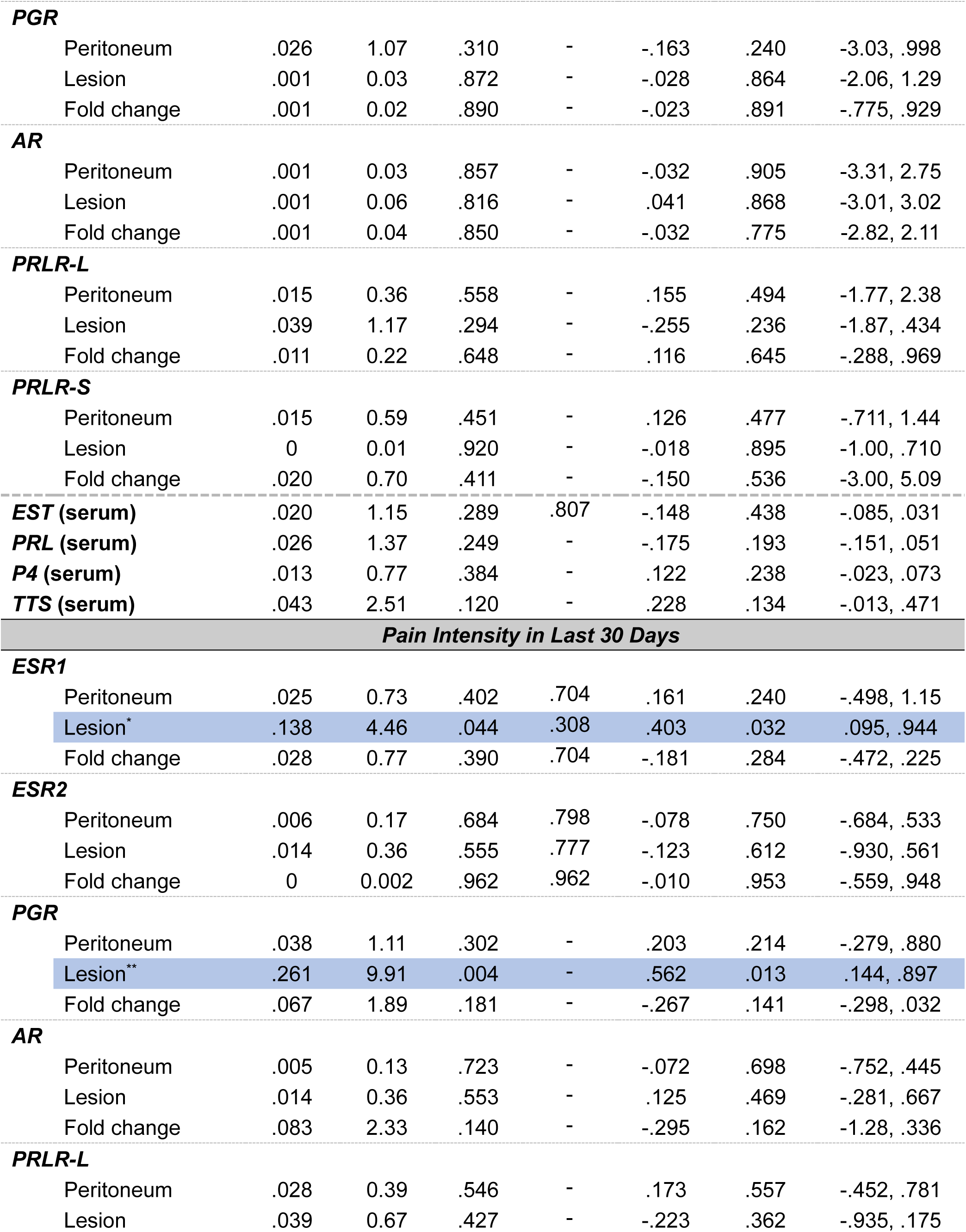
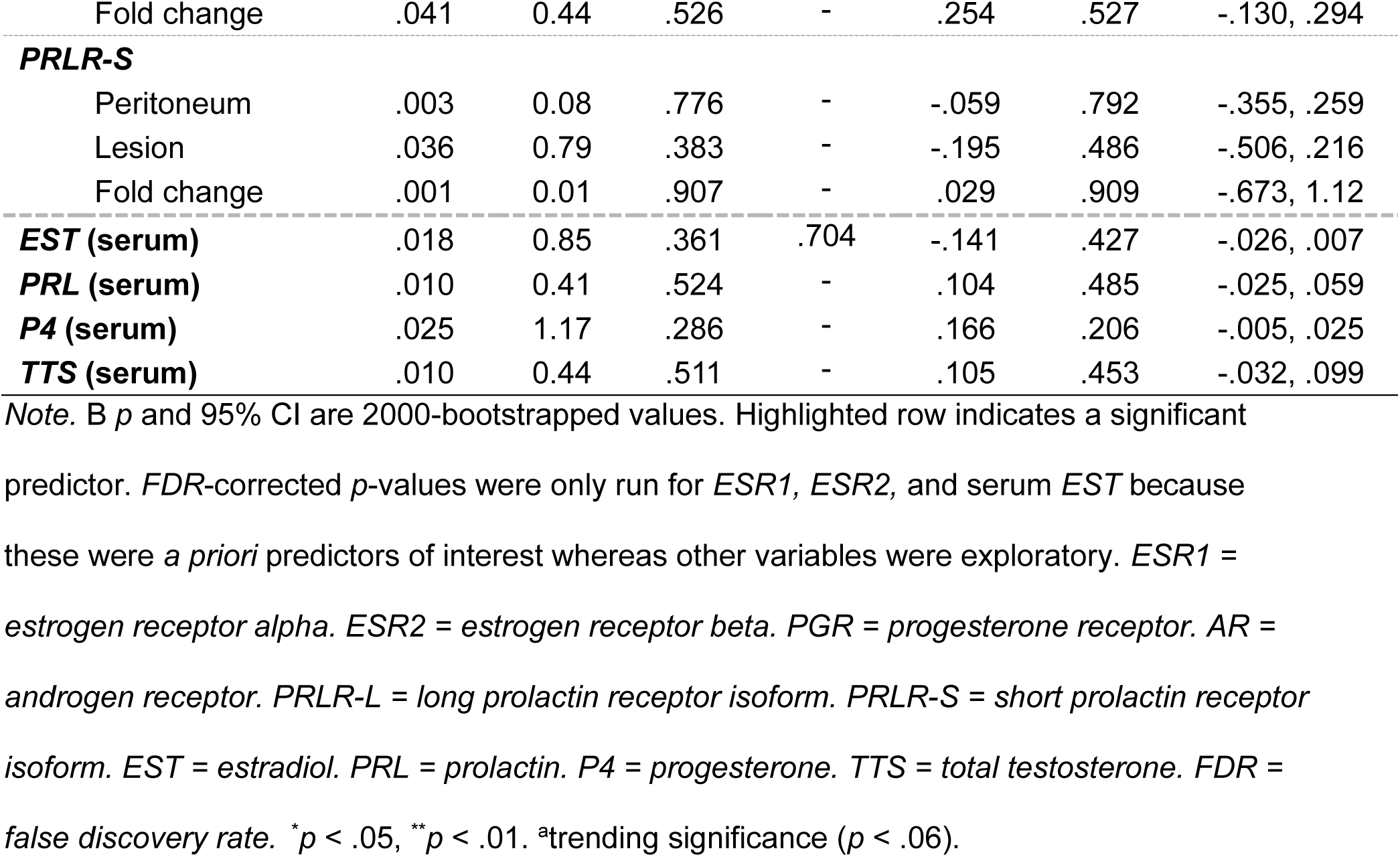
Linear regressions of tissue and serum sex hormones predicting clinical pain outcomes.

